# A multiscale model suggests that a moderately weak inhibition of SARS-CoV-2 replication by type I IFN could accelerate the clearance of the virus

**DOI:** 10.1101/2021.01.25.427896

**Authors:** Anass Bouchnita, Alexey Tokarev, Vitaly Volpert

**Affiliations:** Department of Integrative Biology, University of Texas at Austin, Austin, TX 78712, USA; Peoples Friendship University of Russia (RUDN University) 6 Miklukho-Maklaya St, Moscow, 117198, Russian Federation; Institut Camille Jordan, UMR 5208 CNRS, University Lyon 1, Villeurbanne, 69622, France; INRIA Team Dracula, INRIA Lyon La Doua, Villeurbanne, 69603, France

**Keywords:** COVID-19, SARS-CoV-2, hybrid modelling, multiscale model, immunosenescence, immune response

## Abstract

Severe acute respiratory syndrome coronavirus 2 (SARS-CoV-2) is a highly transmissible RNA virus that emerged in China at the end of 2019 and caused a large global outbreak. The interaction between SARS-CoV-2 and the immune response is complex because it is regulated by various processes taking part at the intracellular, tissue, and host levels. To gain a better understanding of the pathogenesis and progression of COVID-19, we formulate a multiscale model that integrate the main mechanisms which regulate the immune response to SARS-CoV-2 across multiple scales. The model describes the effect of type I interferon on the replication of SARS-CoV-2 inside cells. At the tissue level, we simulate the interactions between infected cells and immune cells using a hybrid agent-based representation. At the same time, we model the dynamics of virus spread and adaptive immune response in the host organism. After model validation, we demonstrate that a moderately weak inhibition of virus replication by type I IFN could elicit a strong adaptive immune response which accelerates the clearance of the virus. Furthermore, numerical simulations suggest that the deficiency of lymphocytes and not dendritic cells could lead to unfavourable outcomes in the elderly population.

## 1 Introduction

Severe acute respiratory syndrome coronavirus 2 SARS-CoV-2 is a novel coronavirus that emerged in Wuhan, China at the end of 2019. The resulting coronavirus disease (COVID-19) has caused a global pandemic that represents a major threat to public health. The response of patients to SARS-CoV-2 infection ranges from no or mild symptoms to moderate and severe symptoms [1]. This intersubject variability suggests that the individual immune response plays a crucial role in determining the clinical evolution of the disease. Indeed, deficiency in CD4 and CD8 T-cells, for example in the elderly population, is associated with unfavourable clinical courses in COVID-19 [2, 3].

The interplay between SARS-CoV-2 and the immune response begins during the early stages of the infection. The internalization of the SARS-CoV-2 is essential for the spread of the virus as it allows the replication of virions. Receptors of SARS-CoV-2 can be found on a wide range of cells in the human body [4]. In addition to epithelial cells, SARS-CoV-2 also replicate inside cells of the immune system such as lung macrophages and dendritic cells (DCs). The infection of dendritic cells plays a central role in the progression of SARS-CoV-2 because these cells are responsible for the production of a wide range of antiviral and proinflammatory cytokines [5] such as type I interferon (IFN-I). A well-timed IFN-I response represents the first line of defence against viral infections and leads to a mild or no symptoms while a delayed response is believed to contribute to the development of severe forms of the disease [6]. Interferons control virus internalization and downregulate its replication in cells [7].

However, SARS-CoV-2 uses several approachs to evade the immune response such as inhibiting IFN signalling [8]. Therefore, numbers game between virus growth and the immune response during the early stages of the infection determines the outcome of the disease [9].

The adaptive immune response starts when antigen-presenting cells (APCs), such as DCs, migrate to lymph nodes (LNs) and trigger the division and maturation of T-cell and B-cell lymphocytes. Following maturation, antigen-specific (Ag-specific) lymphocytes enter the bloodstream and peripheral lymphoid organs where they survey for invading pathogens. Ag-specific CD8 T-cells eliminate infected cells through cell-to-cell contact while Ag-specific B-cell lymphocytes secrete antibodies that eradicate the virus. The production and release of mature T-cells and B-cells require four to five days. Therefore, adaptive immunity only starts after several days of the infection.

Mathematical modelling was applied to gain insights into the kinetics of virus infections and explore the response of patients to potential therapeutics [10]. Systems of ordinary differential equations (ODEs) were formulated to simulate the progression of COVID-19 in patients [11]. The advantage of this approach is that the proposed models can be analyzed analytically to better understand the dynamics of the system [12]. These models can be validated through the comparison of the simulated viral load with the clinically measured one in COVID-19 patients [13]. They can be coupled with pharmacokinetics-pharmacodynamics (PK-PD) models to design efficient and safe treatment regimens [14]. Agent-based modelling is another framework that was applied to study the fine-grained aspects of SARS-CoV-2 spread. A SARS-CoV-2 tissue simulator was proposed to encourage community-driven model development based on an agent-based representation of cells [15]. This simulator relies on a multiscale description for the spread of SARS-CoV-2 in tissues. The kinetics of viral replication inside cells are represented using ODEs, while the spatial distribution of the virus and cytokines were modelled using partial differential equations (PDEs). Another model for SARS-CoV-2 infection used a similar approach and was used to evaluate the effectiveness of treatment protocols [16].

The interplay between SARS-CoV-2 and the immune response is very complex because it involves several processes occurring across multiple levels of physiological organization. The interaction between SARS-CoV-2 and type I IFN at the intracellular level regulates the infection of cells and the replication of the virus, whereas Ag-specific B-cell and T-cell lymphocytes restrict the spread of the virus at the tissue and the organism levels. Hence, a proper description of the dynamics of SARS-CoV-2 infection requires developing new methods of modelling that describe the interplay between the immune response and the virus in more than a single scale. In this work, we address this question by formulating a novel multiscale framework that models SARS-CoV-2 spread in cells, tissues, and organs. After introducing the model, we calibrate some of the unknown parameters to reproduce the kinetics of the viral load observed in clinical samples of a real COVID-19 patient. Next, we use numerical simulations to understand the effect of the early SARS-CoV-2 and type I IFN interplay on the course and progression of the disease. Finally, we apply the model to gain insight into the possible effects of immunological changes caused by immunosenescence on the course of the infection.

## 2 A multiscale model of SARS-CoV-2 infection

We formulate a hybrid discrete-continuous model to describe the immune response to SARS-CoV-2 infection. The model describes the interplay between the virus with the immune response at the intracellular and extracellular levels. It also simulates the evolution of the infection and the magnitude of the immune response at the level of the organism. We have previously used the same approach to describe the dynamics of the immune response in the lymph node during acute infections [17], human immunodeficiency virus (HIV) infection [18], and cancer metastases [18]. The present model simulates the dynamics of SARS-CoV-2 infection in a section of the epithelial tissue and in the respiratory mucosa. At the same time, we describe the production of antigen-specific lymphocytes in the lymph node to estimate the influx of these cells to the infected tissue (Figure 1).

**Figure 1:**
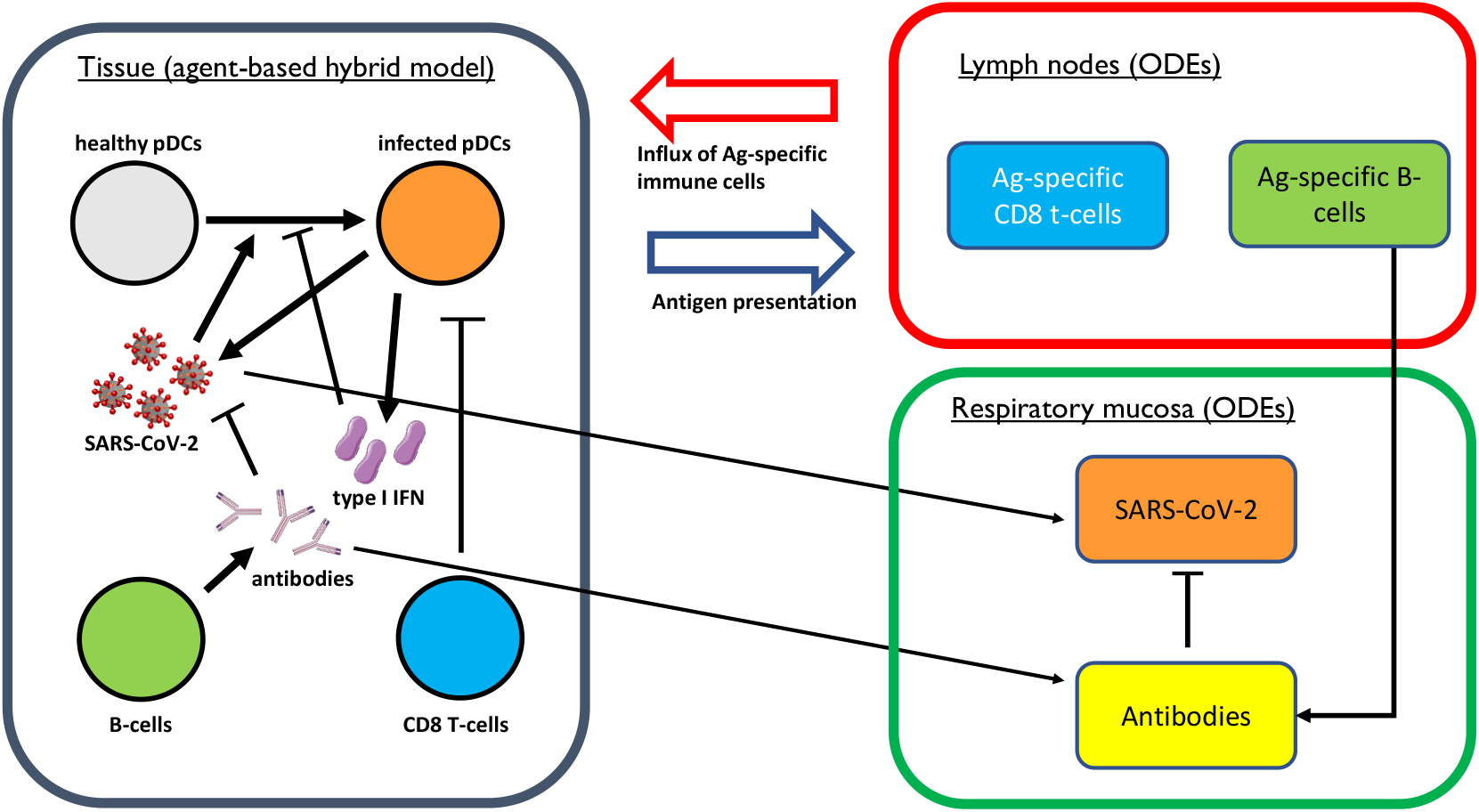
The structure of the model showing the interactions between SARS-CoV-2 and the components of the immune response. The model contains three main compartments: the epithelial tissue, the respiratory tract, and the lymph node. In the tissue, we use an agent-based hybrid description to capture the interactions between cells and represent intracellular and extracellular processes with more accuracy. The second compartments simulates viral kinetics in the upper respiratory tract using a continuous approach. The third adopts the same approach to simulate the production of Ag-specific lymphocytes in the neighbouring lymph nodes. The coupling of the three compartments is ensured by considering that the influx of the virus and antibodies in the respiratory tract depends on their concentration in the epithelial tissue. Furthermore, the influx of immune cell in the tissue depends on their concentration in lymph nodes.

### 2.1 Cell motion

We consider a rectangular computational domain of 1000 *μm* × 1000 *μm* which corresponds to a section of tissue in the upper respiratory tract. Dendritic cells are constantly introduced to the computational domain with a frequency of 2 cells each hour such that the average concentration is maintained at approximately 50 cells. We set the cell cycle to 24 *h* with a random perturbation uniformly sampled in [−2*h*, 2*h*]. Cells move randomly across the domain and we track their motion using Newton’s law of dynamics:

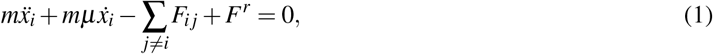

where *m* is the mass of the cell, *μ* is the friction factor, *F^r^* is a random force and *F_ij_* is a repulsive force that describes contact between cells. The modulus of the force *F^r^* is tuned such that the average speed of cells approximate 10 *μm/min*, which is the velocity of T-cells migration in lungs [19].

### 2.2 Intracellular regulation

This virus infects dendritic cells and macrophages and uses them as host for replication. Type I interferon (IFN) plays an important role in the downregulation of virus spread. It induces the expression of several genes (known as interferon-stimulated genes (ISGs)) which downregulate the internalization of SARS-CoV-2 and its replication in host cells [7]. However, SARS-CoV-2 encodes several proteins that allow it to counteract the antiviral activity of IFN. Similarily to SARS-CoV, SARS-CoV-2 was proven to inhibit the induction of interferon [7]. It also inhibits the production of IFN by infected dendritic cells and macrophages [20]. We have summarized the interactions between IFN and SARS-CoV-2 in host cells in Figure 2.

**Figure 2:**
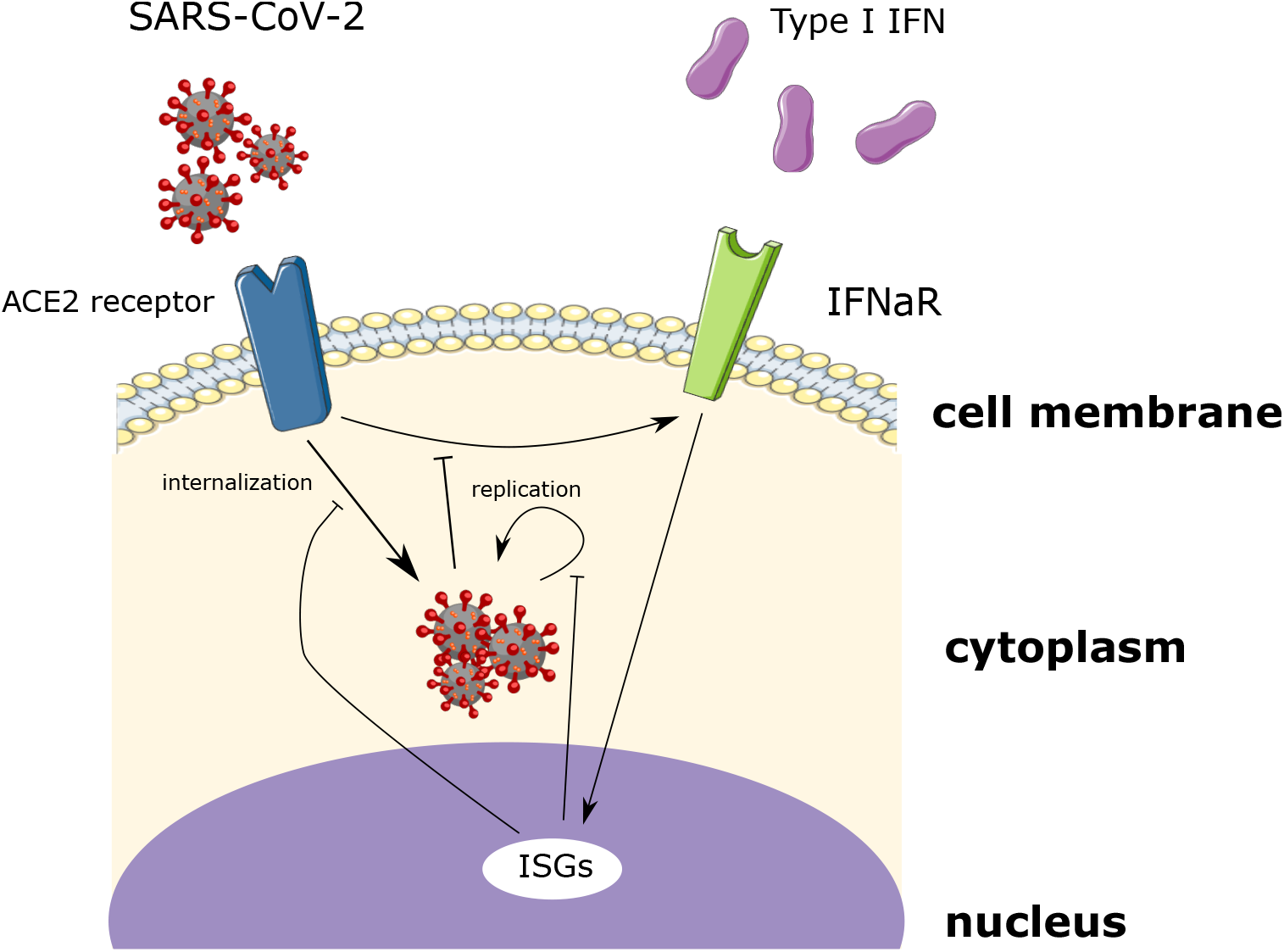
A scheme representing the intracellular interactions between internalized SARS-CoV-2 and interferon-stimulated genes (ISGs) considered in the model.

We use a two-equation model to describe intracellular interaction between interferon and the virus. We start with the equation for internalized SARS-CoV-2 (*v_i_*):

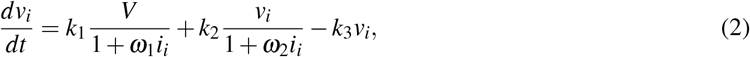

where the first term on the right-hand side of this equation represents the internalization of SARS-CoV-2 and its inhibition by IFN [7], the second term describes the replication of the virius within the cell, and the third term models the degradation of the virus. Next, we describe the concentration of interferon-stimulated genes (ISGs, *i_i_*):

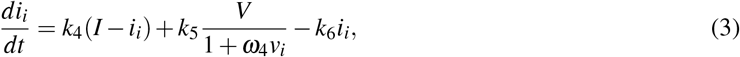

where the first term on the righ-hand side of this equation represents the activation of interferon-stimulated genes through type I IFN and IFNaR interactions, the second describes the activation of ISGs following the induction of IFN receptor by the virus and the inhibition of this process by internalized SARS-CoV-2 [7], while the last represents to the degradation of interferon-stimulated genes.

### 2.3 Extracellular regulation

We describe the extracellular concentration of SARS-CoV-2 and type IIFN. Both of these species are produced by the infected dendritic cells. We begin with the equation for the spatial distribution of SARS-CoV-2:

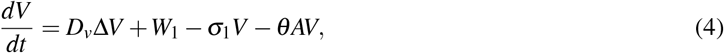

where *D_v_* is the diffusion coefficient of free SARS-CoV-2,and Wi is the production rate of SARS-CoV-2 by the infected DCs:

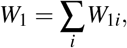

where *W*_1*i*_ = *K*_1_, a positive constant, σ_1_ is the degradation rate constant, and *θ* is the rate constant of virus elimination by antibodies.

Next, we describe the spatial concentration of interferon:

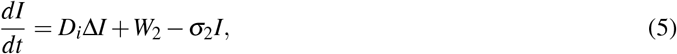

where *D_i_* is the diffusion coefficient of interferon, and *W*_2_ is the production rate which depends on the concentration of internalized virus for each cell:

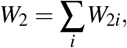

where 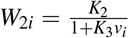, *K*_2_ and *K*_3_ are two positive constants and *v_i_* is the intracellular concentration of the virus. The denominator 1 + *K*_3_*v_i_* represents the inhibition of IFN production by the virus [20], *σ*_2_ the degradation rate constant of type I IFN.

Finally, we describe the concentration of antibodies produced by B-cells:

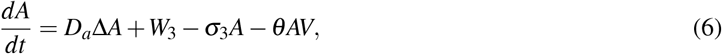

where *W*_3_ = ∑_*i*_*K*_3_, the production term of antibodies by Ag-specific B-cells, the third term on the right-hand side of the equation describes the degradation of antibodies, and the last one represents the consumption of antibodies during the elimination of the virus. We prescribe zero-flux boundary conditions for the three partial differential equations. The initial conditions for the three equation are taken equal to zero.

### 2.4 Dynamics of SARS-CoV-2 infection in the system

#### SARS-CoV-2 viral load

We consider that the influx of infected cells depends on the concentration of the virus in the respiratory mucosa. The cells that enter the epithelial tissue can be either healthy or infected depending on the concentration of the virus in the respiratory tract. We describe the evolution of the total viral load using the following equation:

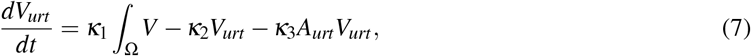

where the first term on the right-hand side of the equation describes the influx of the virus from infected tissue to the respiratory tract, the second term models the decay of the virus, the last term represents the elimination of the virus by antibodies secreted by the B-cells in respiratory airways. We consider that for each introduced cell, the chance that it would be infected is equal to 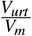, with the highest physiological value possible for the maximal virus load: *V_m_* = 2 × 10^9^ *copies/ml*.

#### Immune cells in the lymph nodes

Adaptive immune response starts approximately after 5 days of the infection. At this moment, we begin the simulation of the concentration of two subtypes of immune cells in lymph nodes: B-cells and CD8 T-cells. Their concentration are described using the following system:

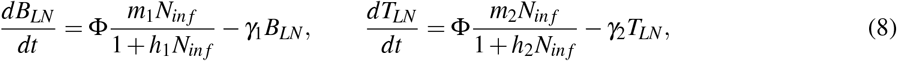

where *B_LN_* and *T_LN_* are the concentrations of B-cells and CD8 T-cells in the lymph node, respectively, *N_inf_* is the number of infected DCs in the tissue. Φ is a patient-specific nondimensional parameter which represents the capacity of the patient to produce Ag-specific lymphocytes. For example, due to changes in the concentration of naïve lymphocytes caused by ageing. We set Φ = 1 as the default value. The concentration of immune cells in the upper respiratory tract determines the influx of these cells to the tissue. B-cells and cytotoxic lymphocytes (CTLs) enter the computational domain depending on their concentration in the lymph node. We consider that each 6 *h*, Ag-specific B-cells and CTLs enter the computational domain with a probability equal to 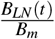 and 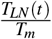, respectively. If the condition of Ag-specific lymphocytes entry to the tissue is satisfied, then the number of cells entering the computational domain will be equal to the absolute value of 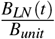 and 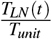, respectively. To keep the number of cells in physiological intervals, we take *B_m_* = *T_m_* = 1500 *cell/μL* and *B_unit_* = *T_unit_* = 100 *cell/μL*. We consider that B-cells secrete antibodies which eliminate the free virus while T-cells eliminate the infected DCs upon direct contact with a given probability *p* equal to 0.28 which was estimated in an experimental study [21].

#### Concentration of antibodies in the respiratory mucosa

The concentration of antibodies in respiratory airways depends on their influx from the tissue and their production by B-cells:

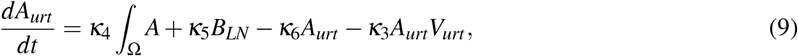

where the first term on the right-hand side of the equation describes the influx of antibodies from the infection site to the respiratory tract, the second term models the release of antibodies by Ag-specific B-cells released from the lymph node, the third corresponds to the degradation of antibodies, and the last describes the consumption of antibodies during the elimination of the virus.

### 2.5 Numerical implementation

We consider a time step of *dt* = 0.02 *min* and a space step equal to *dx* = 10 *μm*. We discretize PDEs using an explicit scheme and we implement them using the finite difference method. The Euler forward method is used to solve ODEs except for the equation of motion which is simulated using an Euler backward scheme. The code was written using C++ in the object-oriented programming (OOP) style. The numerical values of all parameters are provided in the appendix.

## 3 Results

### 3.1 Model validation

We calibrate the values of some parameters to reproduce the kinetics of the viral load in the upper respiratory airways of COVID-19 patients [13]. In this clinical study, virological analysis was conducted for nine COVID-19 patients who showed mild upper respiratory tract symptoms. The viral load reached its maximal value in the first week after the apparition of symptoms in most of these patients. The measurements started after the peak of the viral shedding. For validation purpose, we consider one of these patients who did not have any comorbidities and we tune some of the parameters to reproduce the kinetics of its viral load observed in sputum. In particular, we fit the rates of virus elimination by antibodies in the upper respiratory tract and the infected tissue. Then, we slightly adjust the frequency of CD8 T-cells and B-cells that enter the computational domain such that the slope of the viral load during clearance approximates the observed one (Figure 3, a).

**Figure 3:**
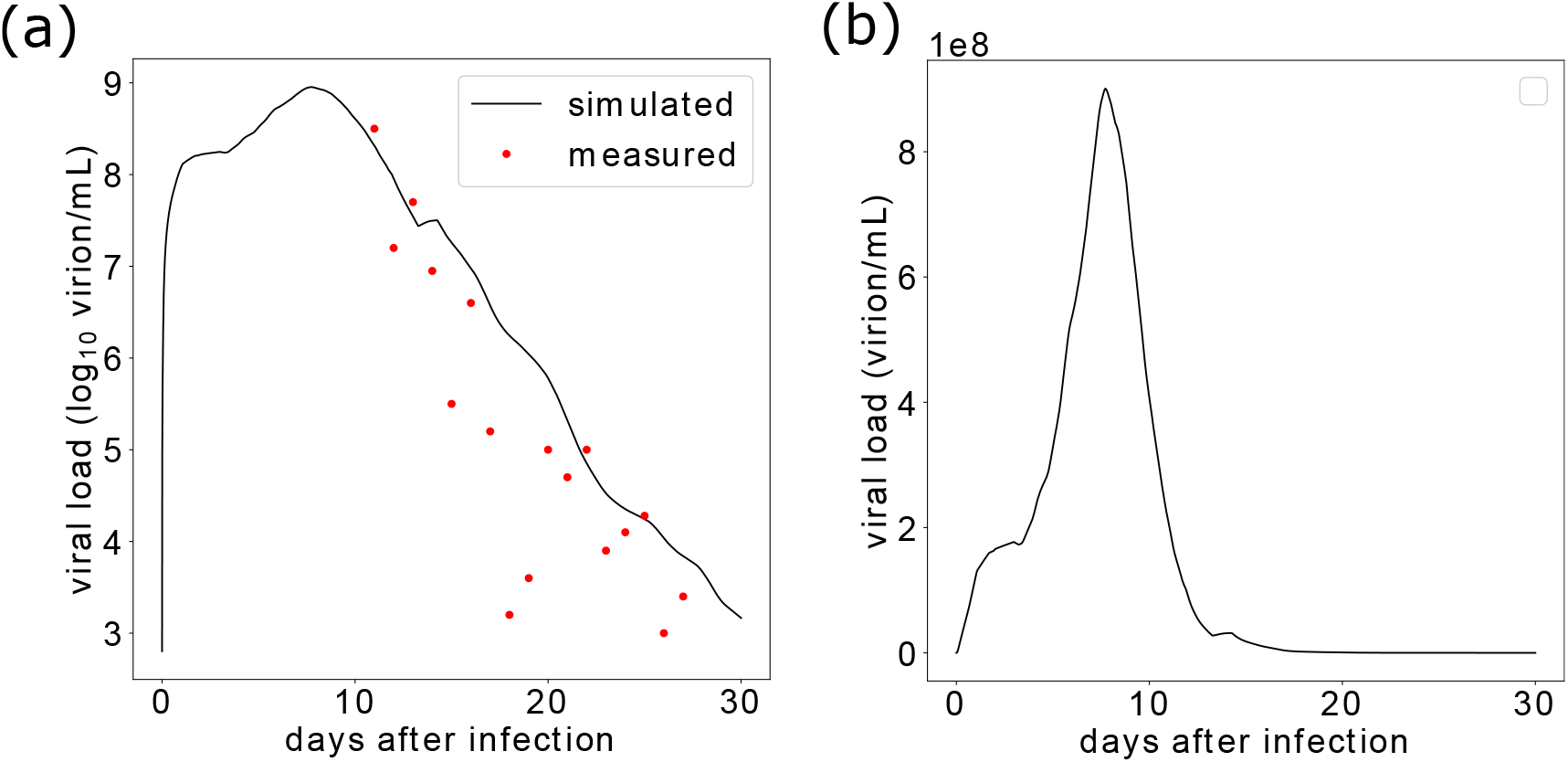
a) Comparison of simulated viral load with clinical data from the sputum sample of a real patient [13] in *log*_10_, assuming an incubation period of 7 days. b) the simulated viral load over time.

The viral load reached a maximal value of 9 × 10^8^ *virion/mL* on day 8 after the infection. Then, it quickly dropped below 1 × 10^8^ *virion/mL* by day 11 as shown in Figure 3, b. The concentration of the virus continues to decrease until reaching a value of 1466 *virion/mL* by day 30. We represent cell dynamics in the tissue in Figure 4. This figure shows the different stages of the infection. In the beginning of the simulation, only a few DCs are infected and produce both the virus and type I IFN. By day 10, the first immune cells are introduced to the computational domain. A few days later, the number of CTLs become sufficiently high to eliminate the majority of infected DCs. By day 25, the majority of CTLs have already died and only B-cells remain. These B-cells produce antibodies which eradicate the free virus. At the end of the simulation, only healthy DCs and a few B-cells remain in the computational domain which marks the end of the infection and the return to the normal state. The evolution of the concentrations of the free virus, type I IFN and antibodies in the tissue are shown in Figure 5.

**Figure 4:**
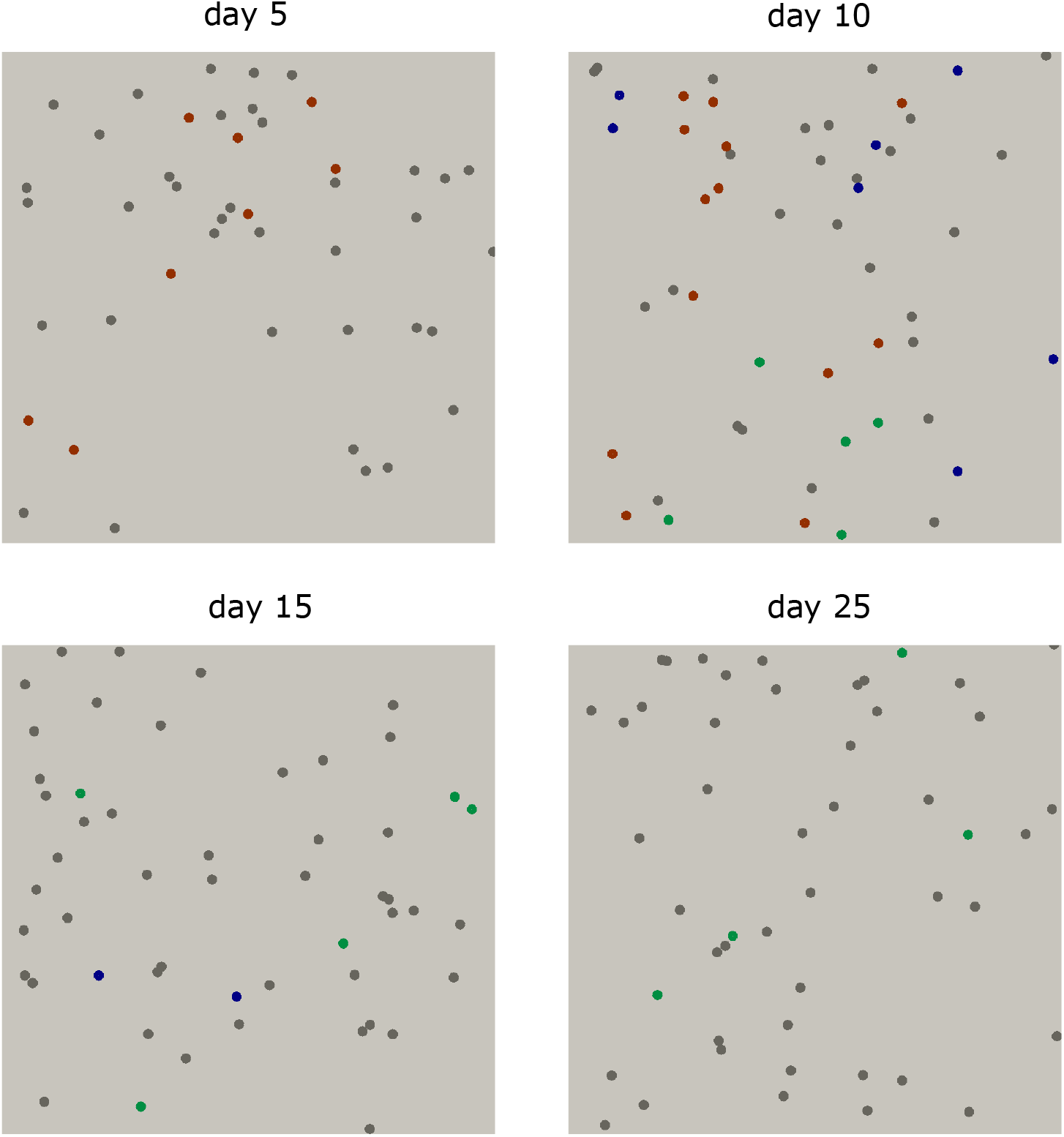
Screenshots of numerical simulations showing the different stages of the infection. Each type of cells is represented with a distinct color: infected DCs are shown in red, healthy DCs in gray, Ag-specific B-cells in green, and Ag-specific CTLs cells in blue. The radius of cells was increased to facilitate visualization.

**Figure 5:**
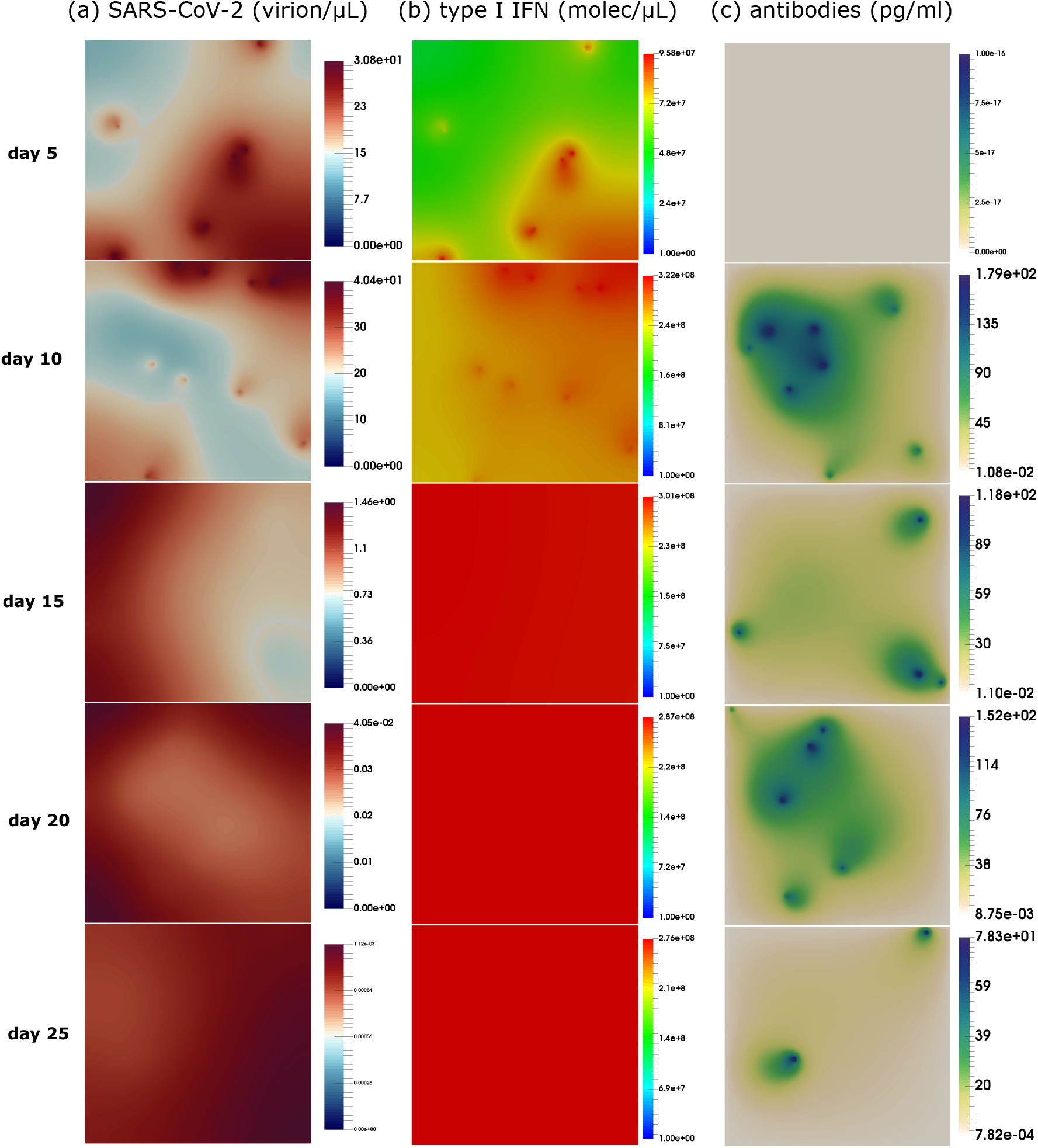
The distribution of the free virus, type I IFN and antibodies at different stages of the infection.

The evolution of the number of cells in the epithelial tissue and the lymph nodes provides a better understanding of the dynamics of SARS-CoV-2 spread. This infection begins with an increase in the number of infected DCs during the first 12 days (Figure 6, a). The number of infected DCs keeps increasing until the arrival of Ag-specific immune cells because the production of free virus keeps getting higher. Immune cells begin entering the computational domain in day 5. The number of CTLs reaches its maximal value in day 12, then it quickly drops to zero by day 20. The number of B-cells reaches its peak and then start slowly decreasing, but with oscillations due to stochasticities of the agent-based model (Figure 6, b). These dynamics are explained by the number of Ag-specific CTLs and B-cells in the lymph nodes (Figure 6, c). A bell-shaped acute CTL response is observed while Ag-specific B-cells remain for a longer time in the lymph nodes. The concentration of antibodies in the respiratory mucosa follows the evolution of B-cells in the epithelial tissue and in lymph nodes (Figure 6, d).

**Figure 6:**
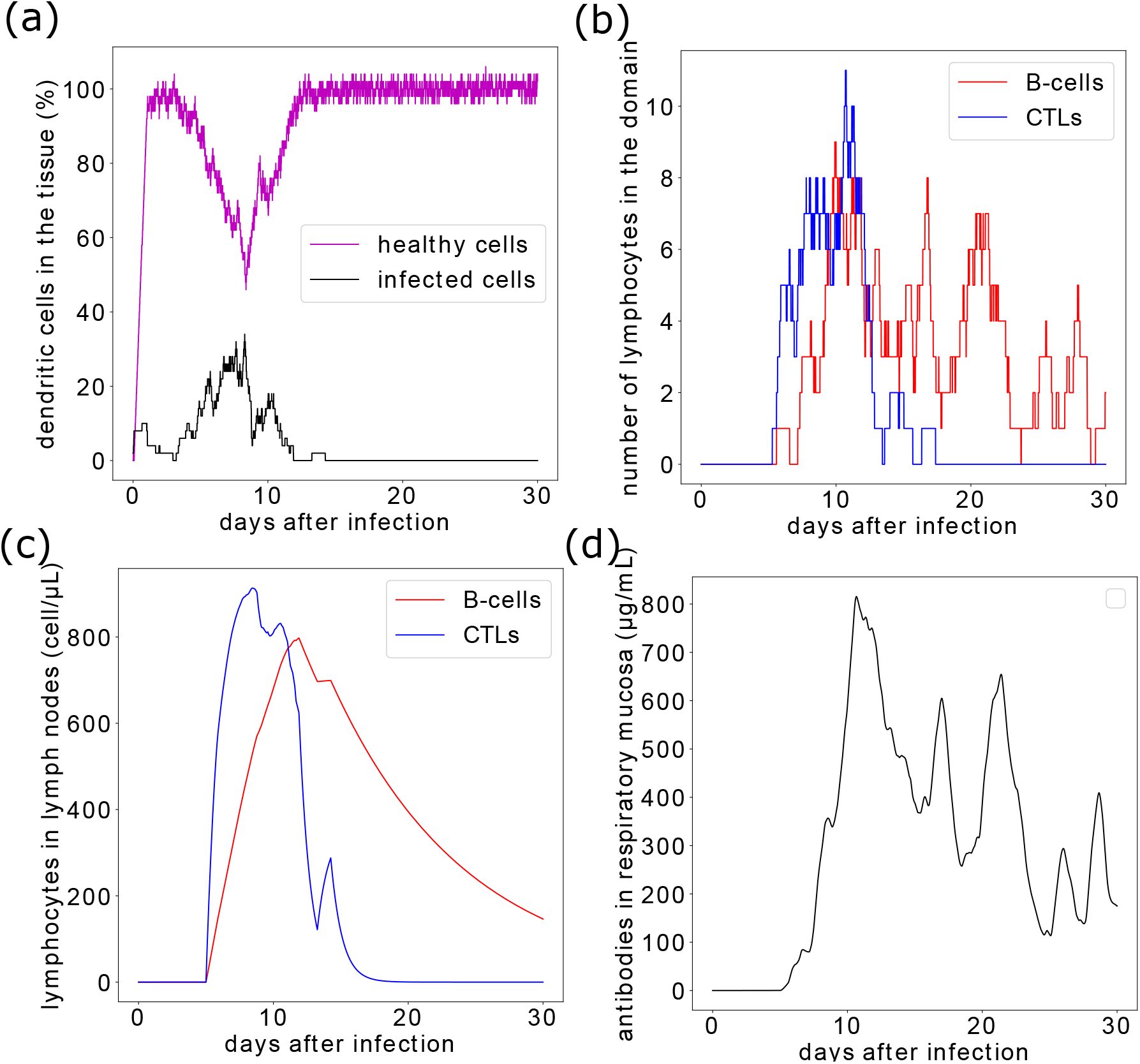
Population of cells and concentration of antibodies in the epithelial tissue, the respiratory tract, and lymph nodes: a) healthy and infected DCs, b) Ag-specific B-cells and CTLs in the epithelial tissue, c) Ag-specific lymphocytes in lymph nodes, d) concentration of antibodies in the respiratory mucosa.

### 3.2 Moderately weak inhibition of virus replication by type I IFN accelerates the clearance of the viral load

We explore the effect of type I IFN on virus replication and how it impacts the clinical course of the infection. To achieve this, we change the value of the parameter ω_2_ in Equation (2) which quantifies the inhibition of virus replication by interferon-stimulated genes. As expected, reducing the value of this parameter increases the number of infected DCs because the replication of the virus inside cells is higher (Figure 7, a). As a result, virus increases production which upregulates the viral load. At the same time, the high number of infected cells increases the magnitude of CD8 T-cells and B-cells production as shown in Figures 7, b and c. The level of produced antibodies also increases as a result of the upregulation in B-cells production (Figure 7, d).

**Figure 7:**
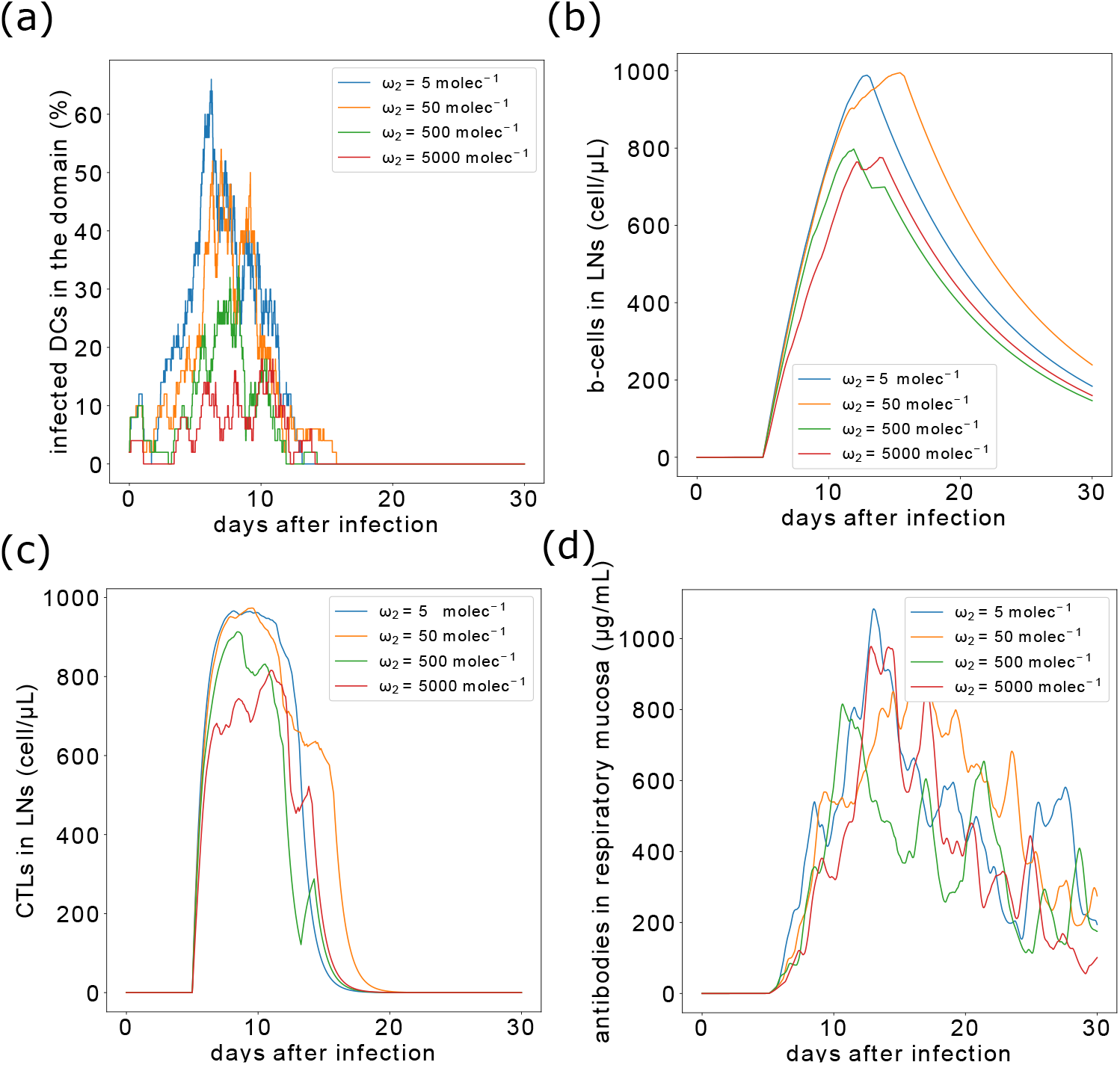
The kinetics of the immune response for three values of the inhibition rates by type I IFN. a) population of infected DCs, b) Ag-specific B-cells in the lymph nodes, c) Ag-specific CTLs, d) concentration of antibodies in the respiratory mucosa.

Next, we study the effect of IFN response on the kinetics of the viral load. The viral load gives an overview of the disease progression and severity. As expected, the shape of the viral load curve does not change but the peak reached by the viral load was significantly higher when the rate of replication inhibition is low (Figure 8, a). However, as we compare the curves of the viral loads in the *log*_10_ scale (Figure 8, it appears that the clearance of the virus is accelerated. For low values of inhibition constants (*ω*_2_ = 5, 50 *molec*^-1^), virus concentration in the respiratory tract falls below the curves of the other values of *ω*_2_ by day 25.

**Figure 8:**
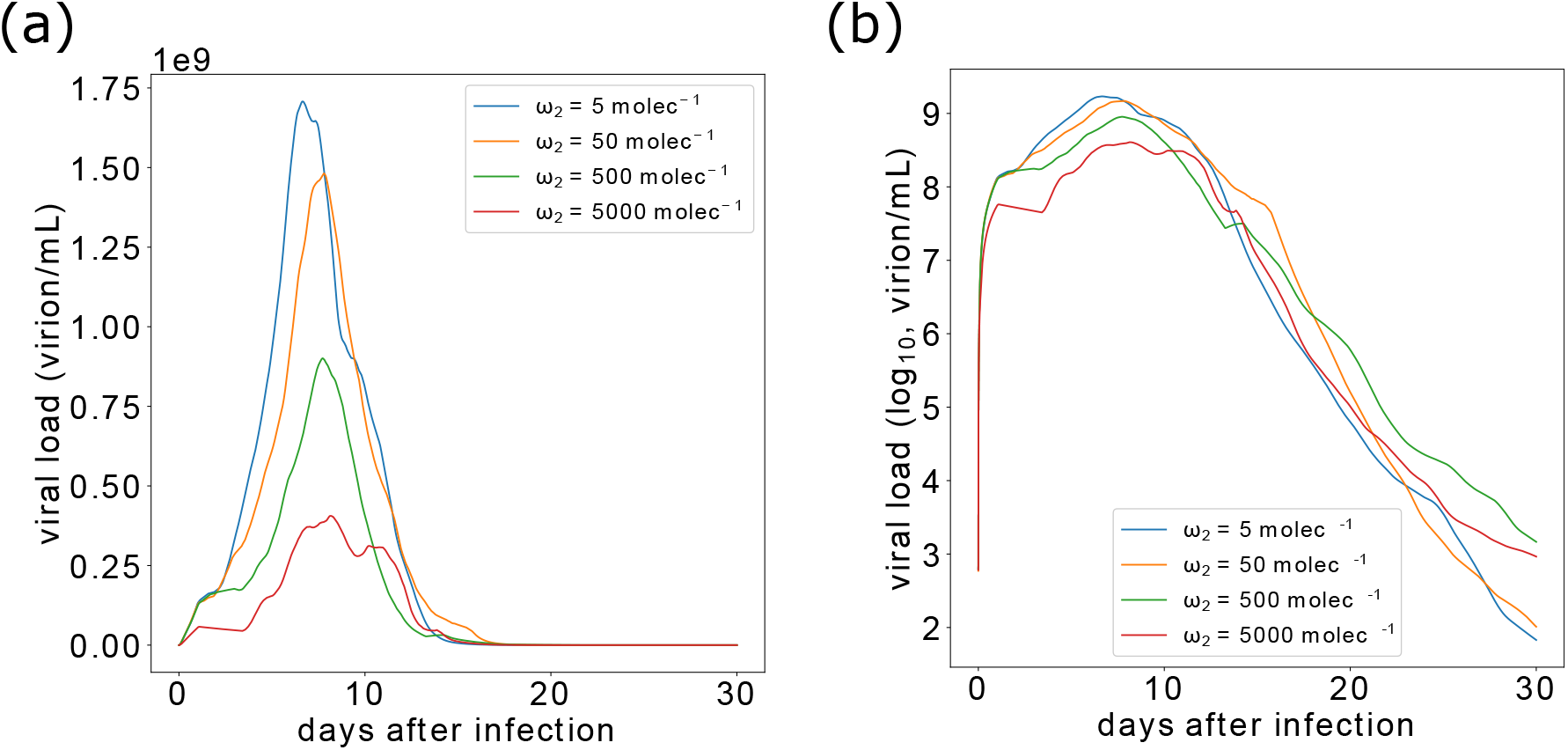
The viral load for three values of the inhibition rates by type I IFN in normal scale (a) and *log*_10_ scale (b). When the inhibition of virus replication is weak, the magnitude of the viral load is higher, but it also decreases faster due to a strong adaptive immune response.

### 3.3 The deficiency of lymphocytes and not DCs extends the duration of the disease in advanced age

Immunosenescence refers to the decline of the immune system due to ageing. The changes occurring during immunosenescence alters the density and function of plasmacytoid DCs. In particular, it was shown that the number of plasmacytoid DCs and their capacity to produce type I IFN decreases with age advancement [22]. DCs play an important role in both the innate and the adaptive immunity and they have several effects on the course and outcome of the disease. First, DCs represent a target for SARS-CoV-2 infection and can promote the replication and spread of the virus. Second, these cells secrete type I IFN which has antiviral action. Finally, they collect antigens and present them to naïve lymphocytes which initiate their differentiation and maturation. Therefore, DCs play opposing roles in the kinetics of SARS-CoV-2 infection. To gain insights into these roles, we added a small perturbation to the number of DCs in the epithelial tissue and study the evolution of the disease.

Decreasing the number of DCs in the computational domain slows down the spread of the disease. As a result, the curve of the viral load is flattened and the virus concentration reached during the peak is reduced (Figure 9, a). However, the flattening of the viral load and the number of infected DCs in the epithelial tissue prolongs the duration of the adaptive immune response which provokes an extended adaptive response and accelerates the clearance of the virus (Figure 9, b). Interestingly, a higher number of dendritic cells amplifies the generation of lymphocytes and antibodies which promotes the clearance of the virus. At the same time, this small elevation in the number of DCs significantly increases the concentration of the virus reached during the peak of the infection. Overall, an increase in the number of dendritic cells accelerate the clearance of the virus. As before, the high infection of DCs elicit a strong adaptive immune response which promotes the clearance of the virus. The dynamics of Ag-specific immune cells in the lymph nodes for the three cases are represented in Figures 9, c and d.

Next, we evaluate the effect of changes in the number of naïve T-cells and B-cells on the clinical course of the infection. Lymphopenia is another hallmark of immunosenescence and believed to be a significant risk factor for severe forms of COVID-19 [23, 24]. To explore the relationship between the count of naïve lymphocytes and the spread of SARS-CoV-2, we alter the value of Φ in Equations (8) which characterizes the capacity of the organism to produce lymphocytes. Figures 10, a and b show that reducing the value of Φ significantly increases the production of the virus and slows down its clearance. The lack of naïve T-cells impairs the ability of the immune system to generate sufficient CTLs and B-cells. As a result, the curves for the evolution of these two cell types in the lymph nodes are flattened as represented in Figures 10, c and d. In the case when the production of lymphocytes is upregulated (Φ = 1.2), the infection is resolved by day 25 as virus concentration falls below 100 *virion/mL*.

**Figure 9:**
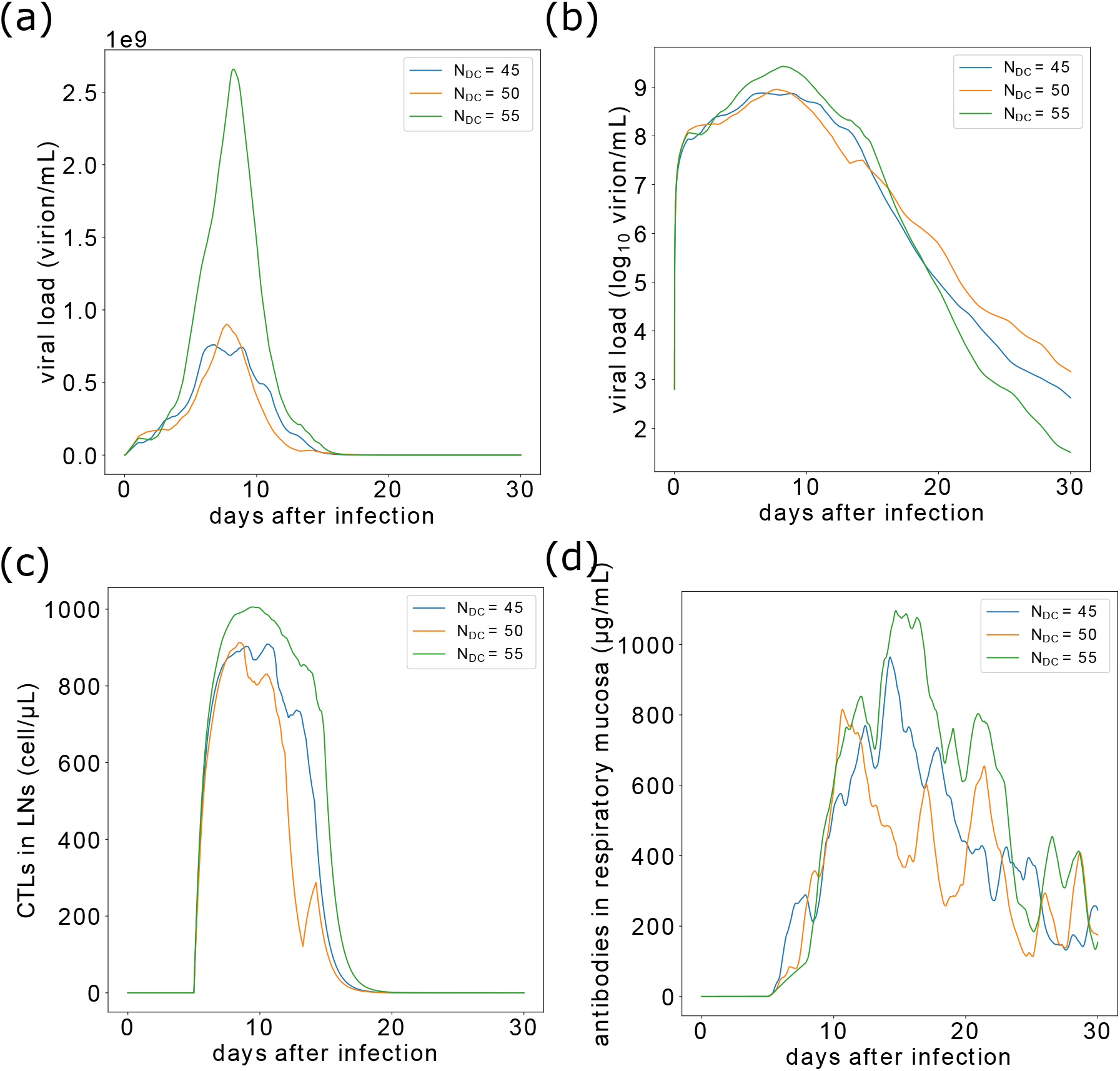
Effects of perturbations in the number of dendritic cells in the epithelial tissue on the evolution of the disease. a) the viral load in normal scale, b) the viral buden in *log_10_* scale. c) the number of Ag-specific b-cells in the lymph nodes, d) the count of CTLs in the lymh nodes.

**Figure 10:**
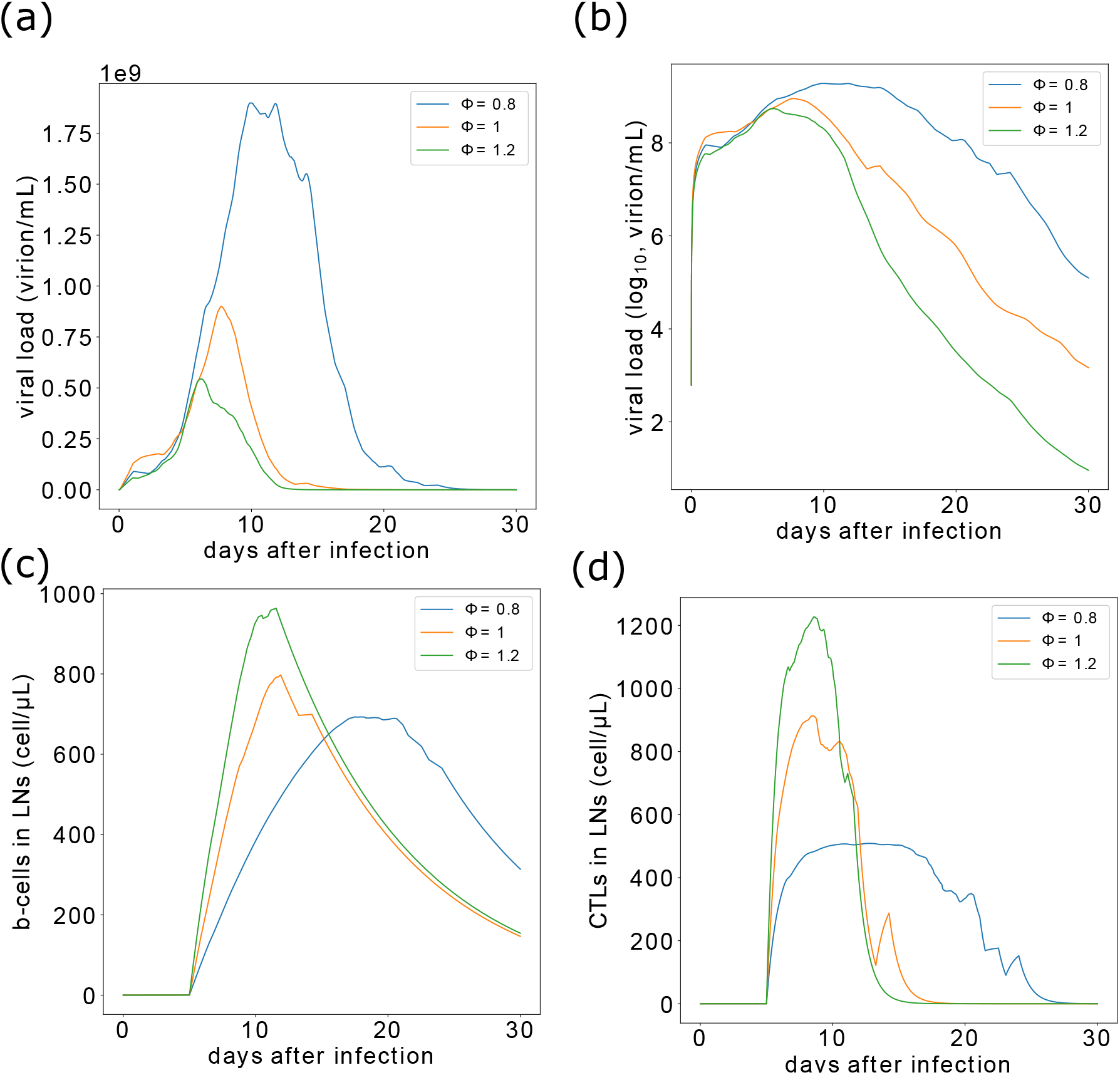
The influence of naïve lymphocyte counts on the evolution of the disease. a) the viral load in normal scale, b) the viral buden in *log*_10_ scale. c) the number of Ag-specific b-cells in lymph nodes, d) the count of Ag-specific CTLs in lymph nodes.

## 4 Discussion

This article studies the interaction between SARS-CoV-2 and the immune response using a multiscale approach. Given the complexity of the problem, we have formulated a new mathematical model encompassing the mechanisms regulating the spread of SARS-CoV-2 across multiple scales. We have used a hybrid discrete-continuous approach to capture the interactions between cells in epithelial tissues with high fidelity. At the same time, we have simulated the kinetics of the viral load in the respiratory tract and the adaptive immune response in the lymph nodes. The coupling between the compartments of the model was ensured by considering that the number of infected DCs in the epithelial tissue affects the production of Ag-specific lymphocytes in lymph nodes, which, in turn, determines the number of lymphocytes the epithelial tissue. The majority of model parameters were taken from the literature. The less known parameters were calibrated such that the simulated viral load approximates the one observed in clinical samples of a real COVID-19 patient [13].

After the validation of the model, we have used it to answer the following question: how interactions between type I IFN and SARS-CoV-2 inside infected cells affects the clinical evolution of the infection? To address the question, we have run several numerical simulations where we have varied the strength of type I IFN in inhibiting SARS-CoV-2 replication. We observed that the weak inhibition of virus replication by type I IFN elicited a strong immune response which accelerated the clearance of the disease. This effect was previously demonstrated for lymphocytic choriomeningitis virus (LCMV) infection in mice through experiments and modelling. It was shown that slow replicating strains of the virus could persist more in the host and evade immune surveillance [25]. It was also previously speculated that slow-growing tumours can “sneak through” immune surveillance [26]. An explanation for this is that the immune response tends to overshoot when faced with acute challenges [27]. The extrapolation of this finding to COVID-19 has several implications. First, it suggests that while a weak and delayed type I IFN response lead to unfavourable outcomes of the disease, a highly strong type I IFN could promote the persisitence of the infection. Second, it indicates that mutations which alter the replication speed of the virus in the new emerging variants could influence not only the severity of the disease but also its duration.

The other important question that we tried to explore concerns the potential effects of ageing on the prognosis of COVID-19. In particular, we aimed to understand what makes immuno-compromised patients more prone to develop severe forms of COVID-19. It was shown in previous studies that ageing reduces the number of DCs and naïve lymphocytes in the organism [22, 23]. According to the predictions of our model, a reduction in DCs does not increase the duration of the disease but can potentially reduce it. This is because DCs represent a target for virus infection and a host for virus replication. Thus, a reduction in their number also reduces the produced viral burden. On the hand, a deficiency in the capacity of the immune system to produce lymphocytes significantly elevates the duration and severity of the infection.

In general, this study demonstrates that the interplay between SARS-CoV-2 and the immune response represents a very complex system with a multiscale organization and nonlinear dynamics. This probably explains the incidence of adverse outcomes in COVID-19. Indeed, small perturbations in the functions of the immune system could provoke significant, and sometimes unexpected, changes in the clinical course of the disease. It is important to note that a few hypotheses were considered while formulating the model. The immune system has a lot of features and peculiarities and it is not possible to include all of them in mathematical models. In our study, we considered only the most relevant components in the immune response to SARS-CoV-2 infection to be able to interpret the obtained results and to reduce the computational cost. Therefore, we did no include natural killer (NK) cells, macrophages, and helper cells. However, their effect is implicitly considered through the choice of parameters. For example, helper cells promote the differentiation of other lymphocytes. Thus, their effect is taken into account through the choice of the production rates for these cells. The pharmacokinetics-pharmacodynamics (PK-PD) of type I IFN-based treatments can be incorporated into the model to determine effective and safe treatment regimens. Furthermore, the model can be used to infer age-dependent parameters for simpler models of the immune response to SARS-CoV-2 infection. These models can be implemented in agent-based models of COVID-19 transmission dynamics [28, 29]. In a forthcoming work, we will incorporate another compartment into our model to simulate the damage caused to the epithelial tissue by the virus and inflammatory cytokines. It was reported that the interactions between epithelial and immune cells correlate with COVID-19 severity [30].

## Acknowledgements

The second and the third authors have been supported by the RUDN University Strategic Academic Leadership Program.

## Appendix: values of model parameters

Most of the numerical values for parameters were taken from the literature. Unknown parameters were fitted to reproduce the observed viral load in clinical samples of a COVID-19 patient [13]. The list of parameters for the models of intracellular and extracellular regulation of the hybrid model are provided in Tables 1, 2, respectively. Table 3 provides the list of parameters used in the model which describe viral kinetics in the respiratory tract and the production of Ag-specific lymphocytes in lymph nodes.

**Table 1:** Numerical values of parameters for the intracellular kinetics between SARS-CoV-2 and type I IFN in dendritic cells.

| Parameter | Value | Description |
| --- | --- | --- |
| $k_1$ | $5.6 \times 10^{-5} \mu L / (virion.h)$ | rate of SARS-CoV-2 internalization [31] |
| $k_2$ | $0.74 \text{ virion}/ml/h$ | rate of SARS-CoV-2 replication [32, 33] |
| $k_3$ | $0.19 \text{ virion}^{-1}$ | rate of SARS-CoV-2 degradation [32, 33] |
| $k_4$ | $2 \times 10^{-4} \text{ molec}/h$ | rate of ISGs activation by IFN [31] |
| $k_5$ | $5 \times 10^{-2} \text{ molec}/(virion.h)$ | induction of IFNaR by type I IFN |
| $k_6$ | $0.1 \text{ molec}^{-1}$ | degradation of IFN |
| $\omega_1$ | $100 \text{ molec}^{-1}$ | inhibition of SARS-CoV-2 internalization by type I IFN. |
| $\omega_2$ | varied | inhibition of SARS-CoV-2 replication by type I IFN. |
| $\omega_3$ | 1 | inhibition of type I IFN by SARS-CoV-2. |

**Table 2:**
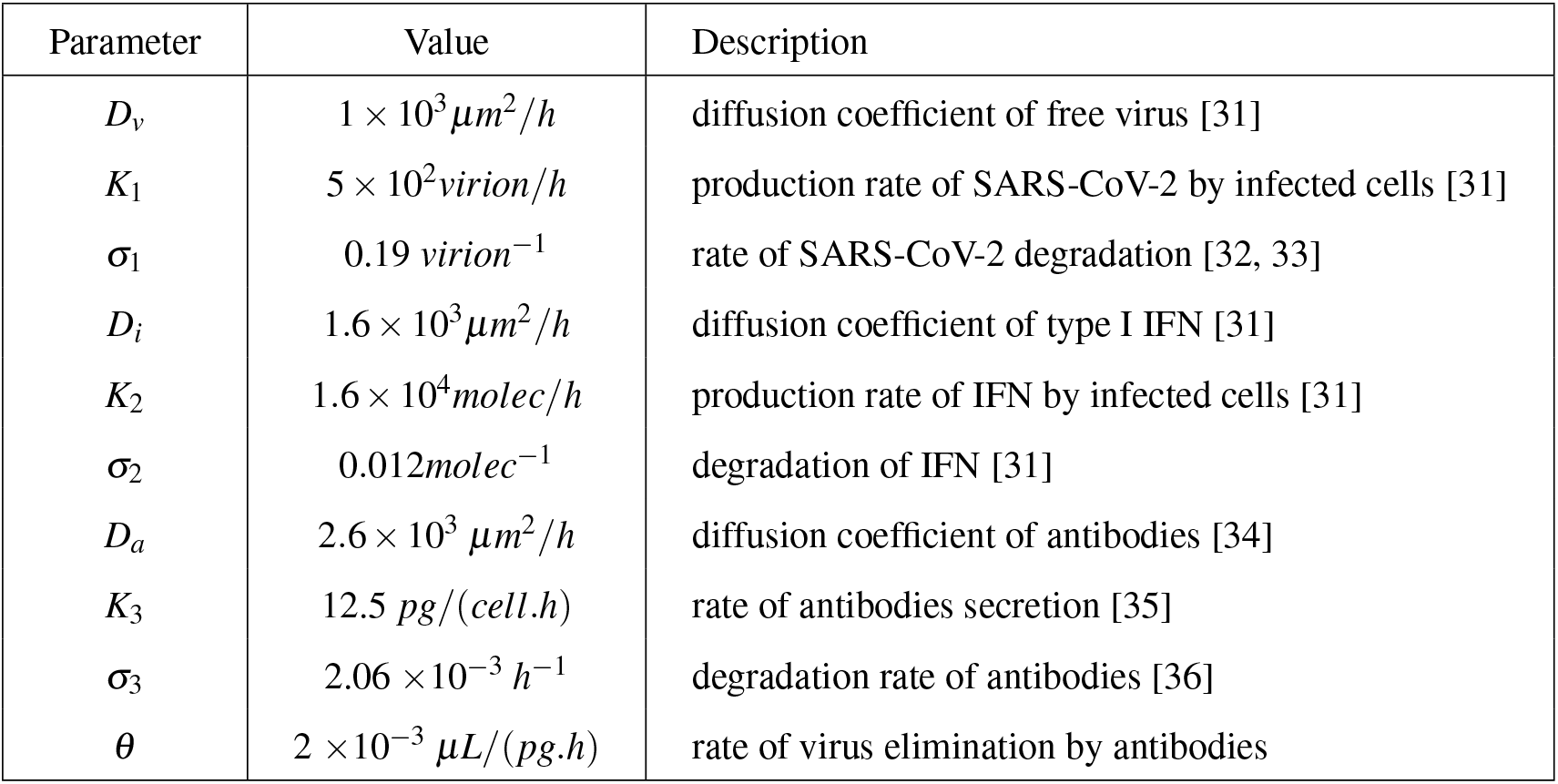
Values of parameters for the extracellular regulation in the model.

| Parameter | Value | Description |
| --- | --- | --- |
| $D_v$ | $1 \times 10^3 \mu m^2/h$ | diffusion coefficient of free virus [31] |
| $K_1$ | $5 \times 10^2 \text{ virion}/h$ | production rate of SARS-CoV-2 by infected cells [31] |
| $\sigma_1$ | $0.19 \text{ virion}^{-1}$ | rate of SARS-CoV-2 degradation [32, 33] |
| $D_i$ | $1.6 \times 10^3 \mu m^2/h$ | diffusion coefficient of type I IFN [31] |
| $K_2$ | $1.6 \times 10^4 \text{ molec}/h$ | production rate of IFN by infected cells [31] |
| $\sigma_2$ | $0.012 \text{ molec}^{-1}$ | degradation of IFN [31] |
| $D_a$ | $2.6 \times 10^3 \mu m^2/h$ | diffusion coefficient of antibodies [34] |
| $K_3$ | $12.5 \text{ pg}/(cell.h)$ | rate of antibodies secretion [35] |
| $\sigma_3$ | $2.06 \times 10^{-3} h^{-1}$ | degradation rate of antibodies [36] |
| $\theta$ | $2 \times 10^{-3} \mu L/(pg.h)$ | rate of virus elimination by antibodies |

**Table 3:** Parameter values for the dynamics of the viral load in respiratory mucosa and of the antigen-specific lymphocytes in the lymph nodes.

| Parameter | Value | Description |
| --- | --- | --- |
| $\kappa_1$ | $1.479 \times 10^3 \mu L/h$ | rate of virus influx to respiratory tract |
| $\kappa_2$ | $3.2 h^{-1}$ | rate of virus degradation [17] |
| $\kappa_3$ | $2.4 \times 10^{-3} h^{-1}$ | rate of virus elimination by antibodies |
| $\kappa_4$ | $9 \times 10^{-2} \mu L/h$ | rate of antibodies influx to respiratory tract |
| $\kappa_5$ | $6 \times 10^{-2} \mu L/(cell.h)$ | rate of antibodies production by B-cells |
| $\kappa_6$ | $2.06 \times 10^{-3} h^{-1}$ | rate of antibodies degradation [36] |
| $m_1$ | $0.45 \mu L^{-1} min^{-1}$ | rate of CTL production |
| $h_1$ | $0.001 cell^{-1} \mu L^{-1}$ | rate of reduction of CTL production due to saturation |
| $\gamma_1$ | $1.25 \times 10^{-2} h^{-1}$ | rate of CTLs natural death [37] |
| $m_2$ | $0.35 \mu L^{-1} min^{-1}$ | rate of Ag-specific B-cells production |
| $h_2$ | $0.001 cell^{-1} \mu L^{-1}$ | rate of reduction of B-cell production due to saturation |
| $\gamma_2$ | $1.25 \times 10^{-3} h^{-1}$ | rate of B-cells natural death [38] |

